# Knowledge, attitudes and biosecurity practices among the small-scale dairy farmers in Sylhet District, Bangladesh

**DOI:** 10.1101/2023.05.28.542608

**Authors:** Tonmoy Chowdhury, Junayed Ahmed, Md Tafazzal Hossain, Mithu Chandra Roy, Md Ashik-Uz-Zaman, Md Nazim Uddin, Md Masudur Rahman, Md Golam Kabir, Ferdaus Mohd Altaf Hossain

## Abstract

**Background:** In the context of zoonosis, Bangladesh’s small-scale dairying is yet to frame satisfactory levels due to poor biosecurity practices.

**Objectives:** This study intended to reveal the degree of knowledge, attitudes, and biosecurity practices among Sylhet district, Bangladesh’s small-scale dairy farmers. We also focused on the association between biosecurity practices and the incidence of non-specific enteritis in humans.

**Methods:** A questionnaire-based survey was conducted on the farmers’ KAP via personal interviews of 15 farmers from the randomly selected fifteen small-scale dairy farms. The questionnaire was developed with six questions for knowledge, six questions for attitude, and 12 questions for the practice of biosecurity measures. Alongside that, data on the number of non-specific enteritis cases experienced by the farmers or their family members were also recorded. Spearman correlation was used to find out the correlation among KAP variables and between practice scores and non-specific enteritis incidences.

**Results:** We found an insignificant (p > 0.05) influence of demographic characteristics over knowledge, attitude, and biosecurity practices. Significant (p<0.05) and strong correlations were found in knowledge-attitude (r = 0.65), knowledge-practice (r = 0.71), and attitude-practice (r = 0.64). Incidences of non-specific enteritis and biosecurity measures’ practice were also strongly correlated (r = -0.9232) and statistically significant (p<0.05).

**Conclusions:** Our study suggests that increasing knowledge and developing a good attitude are necessary to increase the adaptation of biosecurity measures as three of these factors are correlated. Also, farm biosecurity measures are closely related to human health.

## Introduction

The farms act as a source of several pathogenic microorganisms which can cause animal and human health risks (An et al., 2018; Castells & Colina, 2021; Stein & Katz, 2017). Infectious diseases cause severe economic losses to farms as well as result in dissatisfaction among farmers, veterinarians, consumers, and different stakeholders (Makita et al., 2020). In Bangladesh, there is a high risk of infectious disease spread such as Foot and Mouth disease (FMD) (Youssef et al., 2021). Gastroenteritis in humans can also be traced to animal-origin food; for example, enteritis causing *Campylobacter* and *Escherichia coli* (An et al., 2018; Stein & Katz, 2017). To prevent the risk of spreading these types of diseases adaption of biosecurity measures on farms plays an important role (Can & Altuğ, 2014). Adapting good biosecurity measures also helps to improve production efficiency as well (Brennan & Christley, 2012).

However, it is hard to adapt standard biosecurity measures as it depends on various factors like farmers’ knowledge, implementation cost, workforce, implementation complexity, and biosecurity measures differ from region to region (Can & Altuğ, 2014). Before that, in a developing country like Bangladesh, it is important to understand the mindset of the farmers and the factors that influence biosecurity practices which could aid in the implementation of any project regarding biosecurity awareness and practice. There is a lack of studies and available data regarding this topic. Hence, KAP analysis is an efficient tool to draw a conclusion for this purpose. Conducting KAP analysis, it is easier to understand the depth of awareness of the farmers about biosecurity.

Hence, considering above mentioned facts we have conducted the study to understand if the demographic characteristics of the farmers have any influence on biosecurity practice. Also, the nature of association among knowledge, attitude, and practice regarding the biosecurity practice of the farmers. And to find out the association between biosecurity practice and the risk of enteritis in farmers and their family members who are directly or indirectly related to the farms or consume milk from that farm.

## Materials and Methods

### Study Area

This study was conducted on a total of 15 randomly selected dairy farms (farms having not more than 30 animals) in different parts of Sylhet Sadar upazila (24.90568306031467, 91.87500530754328) of Sylhet district; a medium-sized city, situated in the northeast part of Bangladesh (Figure 1). The Sylhet sadar upazila is one of the 13 upazilas under the Sylhet district and has almost every geographical characteristic of all other upazilas including hilly areas and relatively low laying lands as well. It also includes urban and rural area sites as well. So, the farms from Sylhet sadar upazilas that were included in the study will show almost a similar image of the Sylhet district.

**Figure 1:**
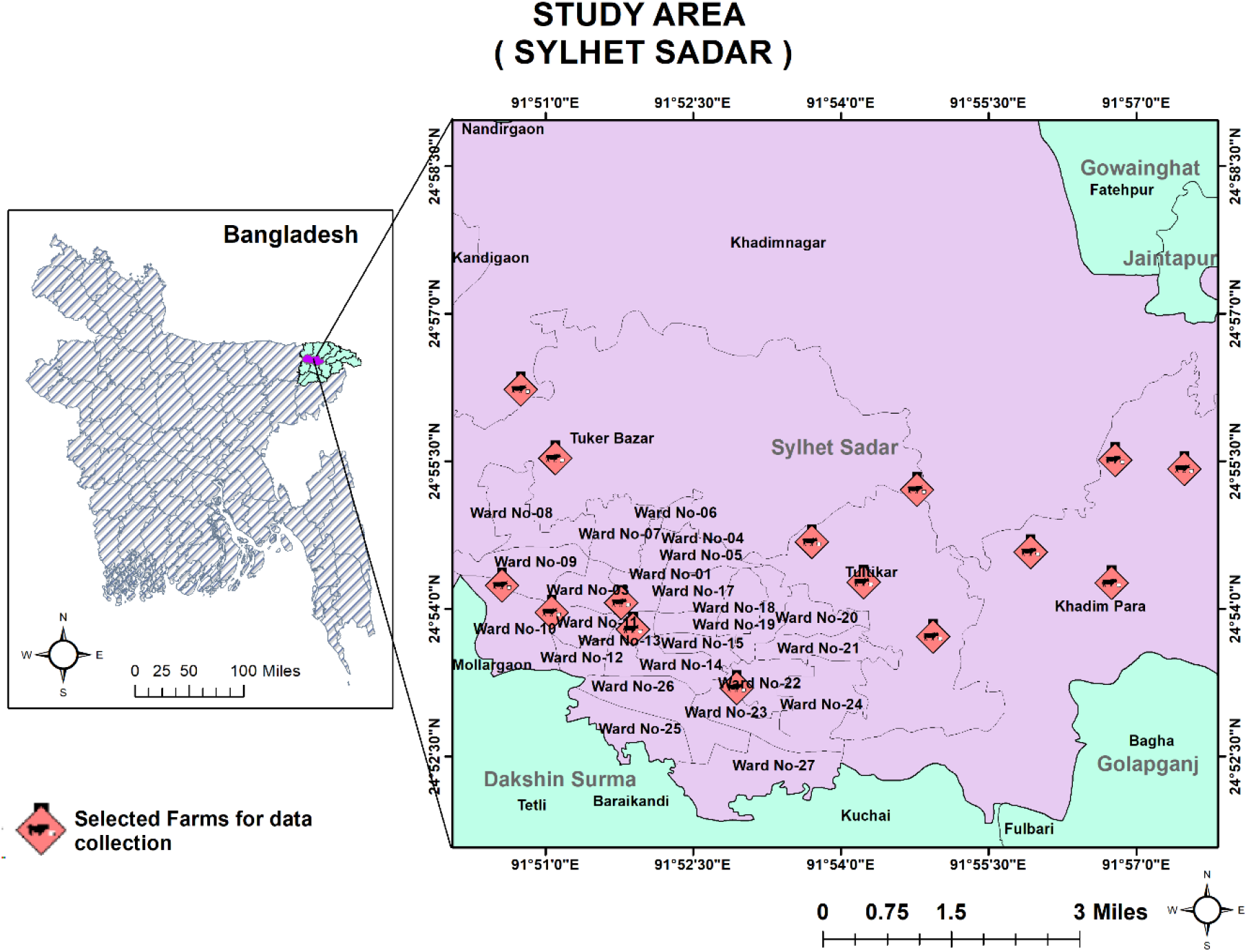
Map of the study area (Sylhet Sadar upazila, Sylhet district, Bangladesh).

### Data collection

Unfortunately, there was no reliable official data available regarding the number of small-scale dairy farms. However, we were able to locate 37 small-scale dairy farms in the Sylhet Sadar region that were operating and actively delivering their dairy products to market, and 23 of the farms agreed to participate in the interview. Out of those 23 farms, we randomly chose 15 farms to ensure that there was no bias and to ensure random selection. Using the prescribed questionnaire, we collected the related data by personally interviewing the farmers (15 farmers; one from each farm) from the 15 randomly selected farms in January 2022 and recorded on Microsoft excel 2021. Knowledge, attitude, and practices regarding biosecurity may vary based on different regions; as a result, the questionnaire was developed by modifying the question sets from two previous studies conducted in Japan and Turkey (Can & Altuğ, 2014; Makita et al., 2020). The questionnaire had a total of 30 questions and we divided the questions into 4 sectors- (1) Demographic characteristics (6 questions; D1 to D6), (2) Knowledge (6 questions; K1 to K6), (3) Attitude (6 questions; A1 to A6) and (4) Practice (12 questions; P1 to P12). For knowledge, attitude, and practice, we set two choices to answer a question- ‘Yes’ and ‘No’. For each positive response (Yes) the responder was given one point and for a negative response (‘No’) no point was rewarded. The possible lowest scores for knowledge, attitude, and practice could be zero (0) and the possible highest scores for knowledge, attitude, and practice could be 6, 6, and 12 respectively.

The data about the incidence of non-specific enteritis (unknown etiology) experienced in the last 2 months by farmers or their family members who either consume the farm milk or work on the farm were collected along with the above-mentioned questionnaire. If the individual experienced diarrhea (loose stool) more than 3 times in 24 hour period with or without other additional symptoms like abdominal pain, nausea, and mucous in stool was considered non-specific enteritis (Baqui et al., 1991; Dey et al., 2007). However, if the individual was having any other illness or medication that could develop diarrhea or other additional symptoms (abdominal pain, nausea, and mucous in stool) was not included in the non-specific enteritis record. Furthermore, we recorded only those cases as non-specific enteritis in which the patient had to seek medical attention and enteritis were diagnosed by a registered clinician.

### Statistical analysis

We did descriptive analysis to find out the frequency, mean, and standard deviation (SD) of the variables. The scores of different variables (Knowledge, Attitude, and Practice) were treated as continuous variables. Test of normality was also performed to identify the distribution of the data. Then we performed non-parametric Independent-Samples Kruskal-Wallis test to determine the association among different variables such as demographic characteristics, knowledge scores, attitude scores, practice scores, etc. We used IBM SPSS Statistics v.26.0.0.0 for that statistical analysis. Finally, we conducted Spearman’s correlation test among scores of knowledge, attitude, and practice using GraphPad Prism 9.3.1 to determine the correlation coefficient (r). Spearman’s correlation test was also conducted to determine the correlation between biosecurity practice score and non-specific enteritis incidence. The significance levels of all the tests were p < 0.05.

## Results

### Frequency percentages and mean scores of knowledge, attitude and practice

The frequency percentages and mean scores for individual questions of knowledge (K1 to K6) and attitude (A1 to A6) are shown in (Table 1). The highest positive response (80%) was found in K4 and the lowest (46.7%) was found in K2 and K6. The frequency percentage of K1, K3, and K5 were equal (60%) (Table 1). In the case of AS, the highest positive response was found in A5 (73.3%) and the lowest was in A2 (33.3%) (Table 1). The percentage of positive response of A1, A3, A4, and A6 was 53.3%, 60%, 53.3%, and 60% respectively (Table 1). The frequency percentages and mean PS (P1 to P12) are shown in (Table 2). The highest positive response (93.3%) was found in P6 and P10 and the lowest (13.3%) was found in P11 (Table 2). P1 and P7 showed the second-highest positive response (86.7%) (Table 2).

**Table 1:**
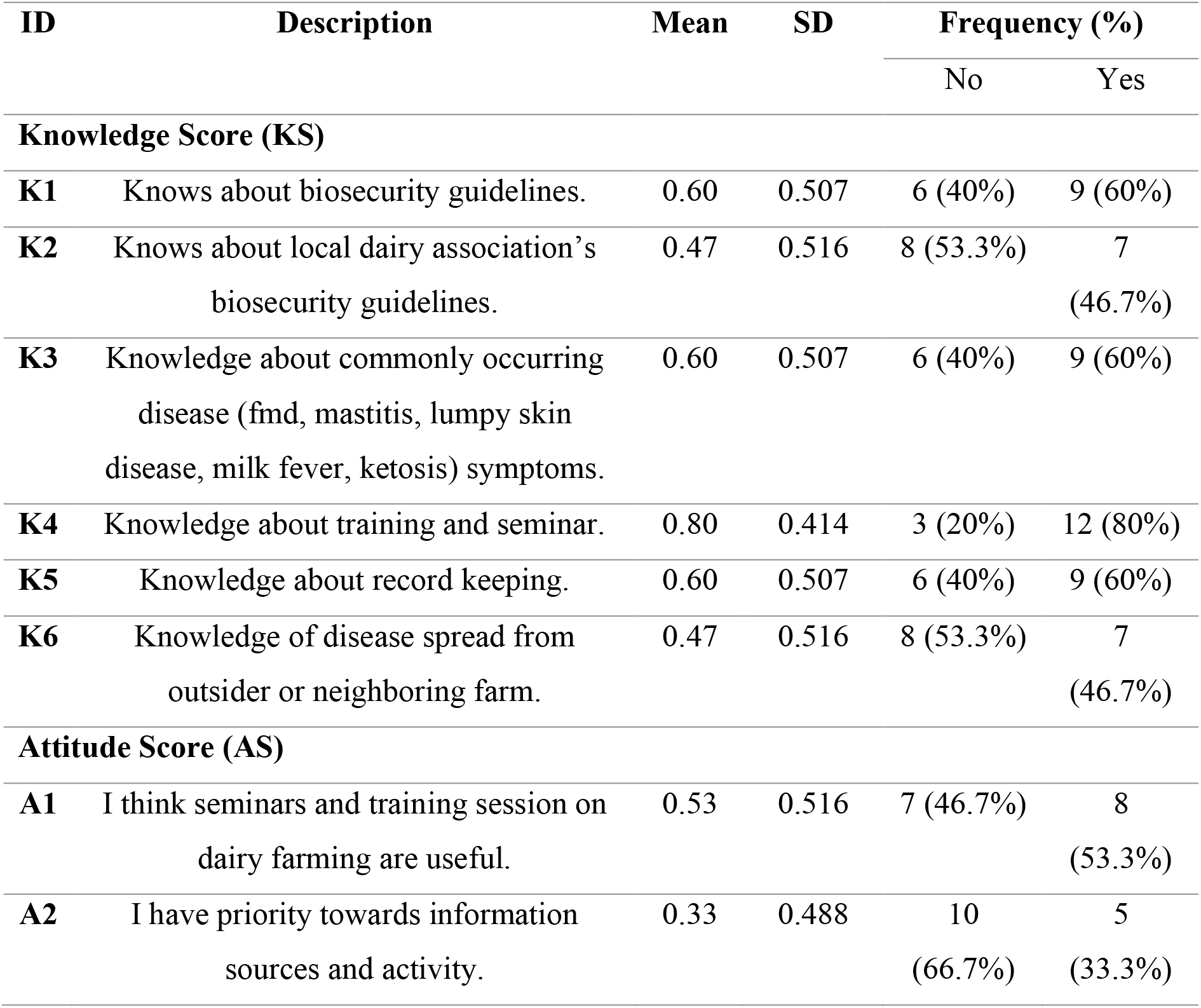
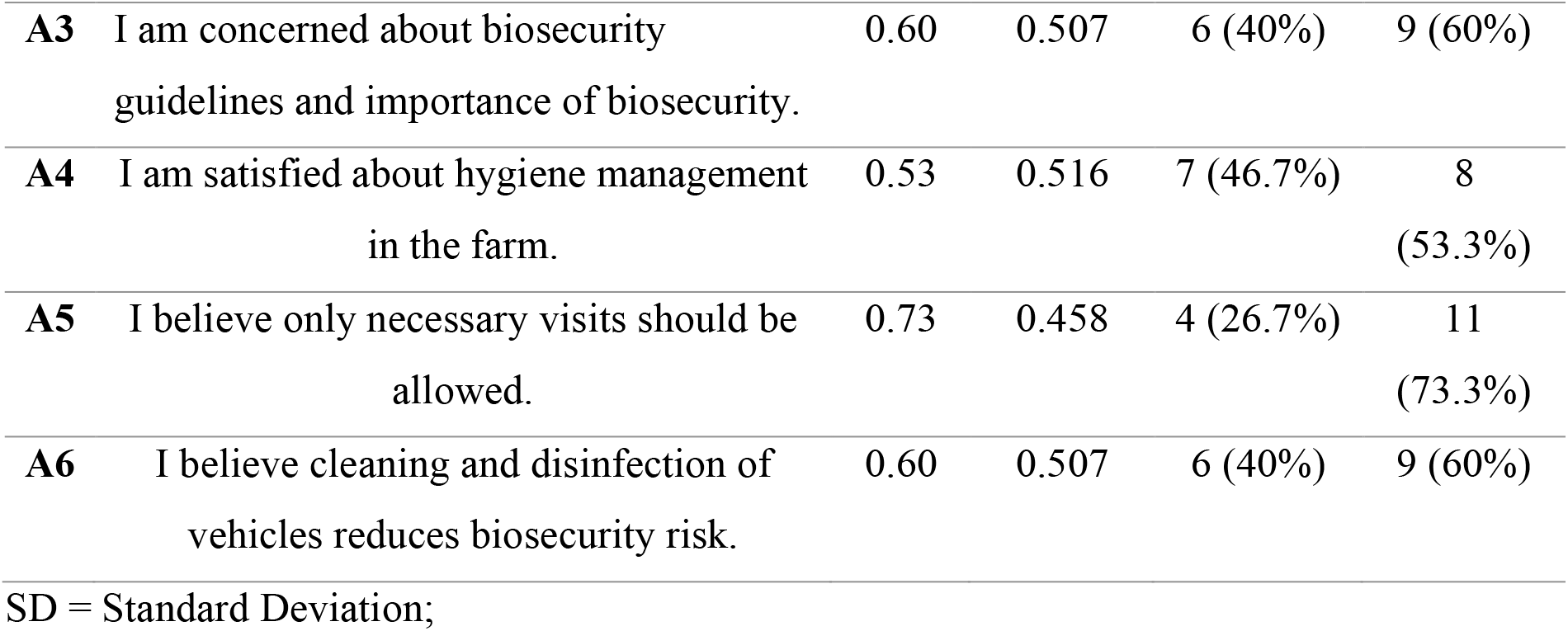
Knowledge and attitude scores of the farmers (N=15) regarding farm biosecurity.

| ID | Description | Mean | SD | Frequency (%) |  |
| --- | --- | --- | --- | --- | --- |
|  |  |  |  | No | Yes |
| Knowledge Score (KS) |  |  |  |  |  |
| K1 | Knows about biosecurity guidelines. | 0.60 | 0.507 | 6 (40%) | 9 (60%) |
| K2 | Knows about local dairy association’s biosecurity guidelines. | 0.47 | 0.516 | 8 (53.3%) | 7 (46.7%) |
| K3 | Knowledge about commonly occurring disease (fmd, mastitis, lumpy skin disease, milk fever, ketosis) symptoms. | 0.60 | 0.507 | 6 (40%) | 9 (60%) |
| K4 | Knowledge about training and seminar. | 0.80 | 0.414 | 3 (20%) | 12 (80%) |
| K5 | Knowledge about record keeping. | 0.60 | 0.507 | 6 (40%) | 9 (60%) |
| K6 | Knowledge of disease spread from outsider or neighboring farm. | 0.47 | 0.516 | 8 (53.3%) | 7 (46.7%) |
| Attitude Score (AS) |  |  |  |  |  |
| A1 | I think seminars and training session on dairy farming are useful. | 0.53 | 0.516 | 7 (46.7%) | 8 (53.3%) |
| A2 | I have priority towards information sources and activity. | 0.33 | 0.488 | 10 (66.7%) | 5 (33.3%) |
| **A3** | I am concerned about biosecurity guidelines and importance of biosecurity. | 0.60 | 0.507 | 6 (40%) | 9 (60%) |
| **A4** | I am satisfied about hygiene management in the farm. | 0.53 | 0.516 | 7 (46.7%) | 8 (53.3%) |
| **A5** | I believe only necessary visits should be allowed. | 0.73 | 0.458 | 4 (26.7%) | 11 (73.3%) |
| **A6** | I believe cleaning and disinfection of vehicles reduces biosecurity risk. | 0.60 | 0.507 | 6 (40%) | 9 (60%) |

SD = Standard Deviation;

**Table 2:** Practice scores (PS) of the farmers (N=15) regarding farm biosecurity.

| ID | Description | Mean | SD | Frequency (%) |  |
| --- | --- | --- | --- | --- | --- |
|  |  |  |  | No | Yes |
| **P1** | Test diseases before buying. | 0.87 | 0.352 | 2 (13.3%) | 13 (86.7%) |
| **P2** | Quarantine for new animal on arrival. | 0.53 | 0.516 | 7 (46.7%) | 8 (53.3%) |
| **P3** | Inspection made by veterinarian on arrival of new animal. | 0.53 | 0.516 | 7 (46.7%) | 8 (53.3%) |
| **P4** | Use hygiene precautions before handling animal feed. | 0.80 | 0.414 | 3 (20%) | 12 (80%) |
| **P5** | Isolating Sick animals. | 0.40 | 0.507 | 9 (60%) | 6 (40%) |
| **P6** | Treatment of Sick animals until clinical sign disappear. | 0.93 | 0.258 | 1 (6.7%) | 14 (93.3%) |
| **P7** | Vaccination against common contagious diseases. | 0.87 | 0.352 | 2 (13.3%) | 13 (86.7%) |
| **P8** | Culling animals that are unresponsive to treatment. | 0.80 | 0.414 | 3 (20%) | 12 (80%) |
| **P9** | Having insect and rodent control plan. | 0.33 | 0.488 | 10 (66.7%) | 5 (33.3%) |
| **P10** | Regular Cleaning and Disinfection of farm. | 0.93 | 0.258 | 1 (6.7%) | 14 (93.3%) |
| **P11** | Footbath in the entrance. | 0.13 | 0.352 | 13<br>(86.7%) | 2<br>(13.3%) |
| **P12** | Clean udder before and after milking. | 0.80 | 0.414 | 3 (20%) | 12 (80%) |

SD = Standard Deviation;

### Comparison of knowledge, attitude and practice scores of demographic characteristics

Comparison of the mean knowledge score (KS), mean attitude score (AS) and mean practice score (PS) according to demographic characteristics are shown in (Table 3). For D1, we found the highest mean KS (4.0) and mean PS (9.3) in the>40 years age group, and the highest mean AS (3.7) in the < 30 years age group (Table 3). For D2, the highest mean KS (3.7) and mean AS (3.8) were observed in the Secondary education group, whereas the highest mean PS (8.6) was identified in the Graduation group (Table 3). For D3, we identified the highest mean KS (3.7) in the group with less than 10 years of farming experience, as well as the highest mean AS (4.5) and mean PS (8.5) in the group with more than 20 years of farming experience (Table3). For D4, there was no responder in the income group of less than $250, the highest mean KS (3.7) was discovered in the income group of $250-500 per month, and the highest mean AS (3.6) and mean PS (8.1) were found in the income group of more than $500 per month (Table3). In D5, the 3–5-year farm’s age group had the highest mean KS (4.5), mean AS (4), and mean PS (8.6) (Table 3). For D6, the highest mean KS (4.3) and mean AS (4) were recorded in the group of fewer than 15 animals on the farm, whereas the highest mean PS (8.6) was identified in the group of more than 25 animals in the farm (Table3). However, the differences between KS, AS, and PS among demographic characteristics (D1 to D6) were insignificant (p > 0.05) (Table 3).

**Table 3:**
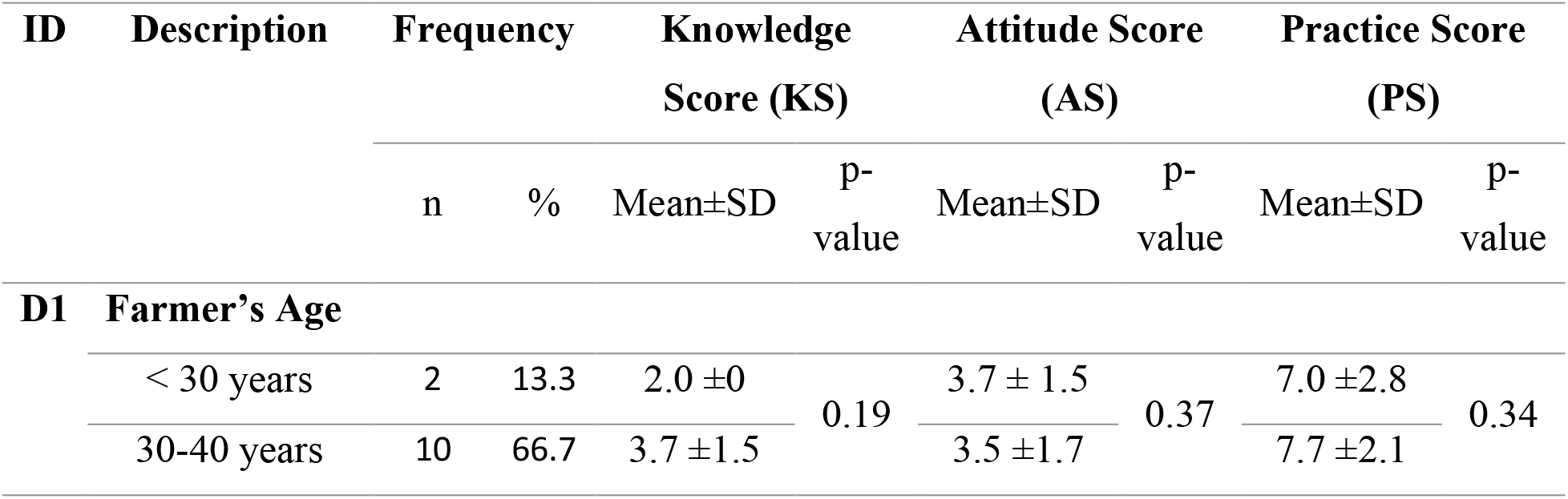
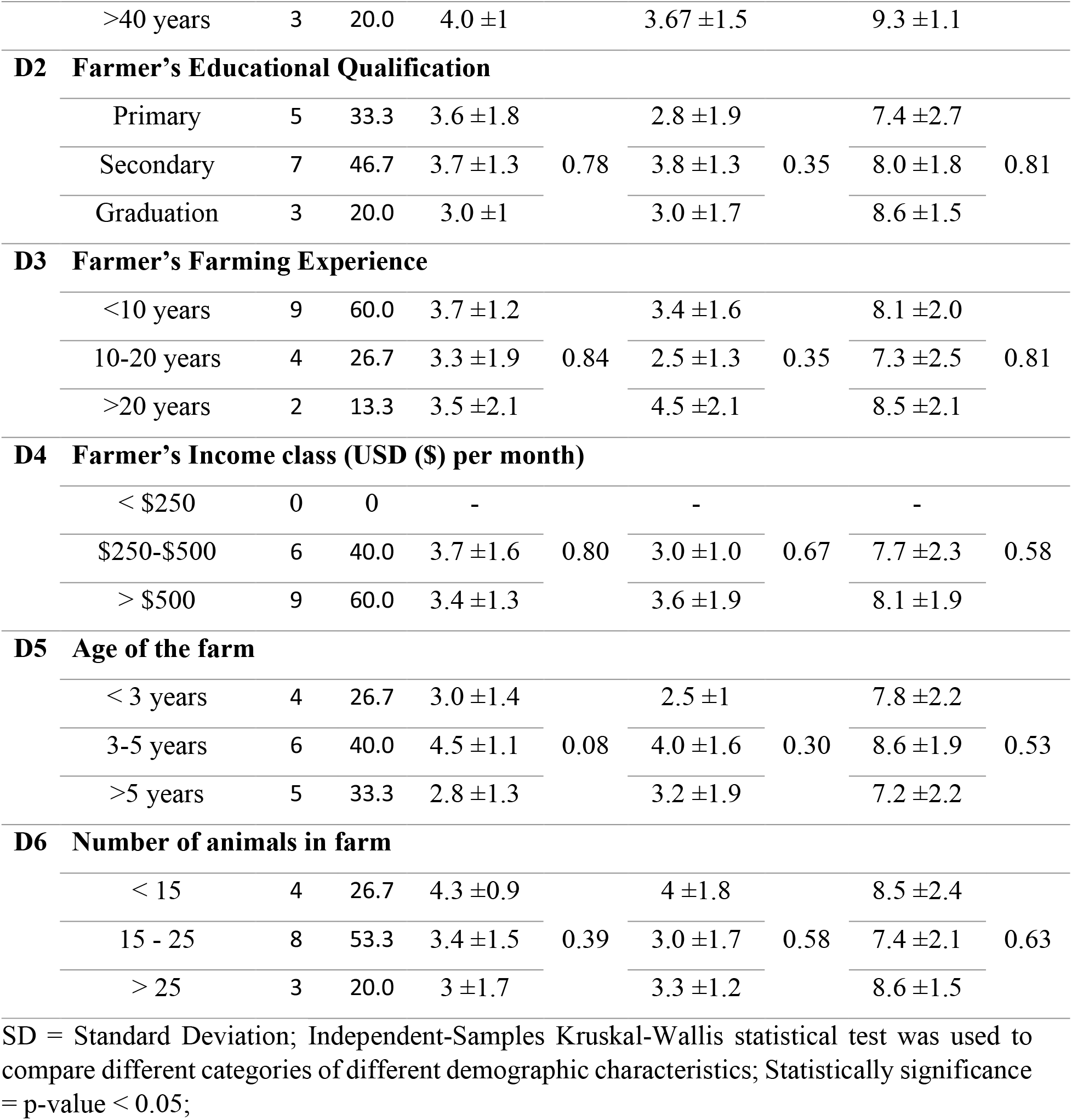
Impact of demographic characteristics on knowledge, attitude and biosecurity practice measures of the farmers (N=15).

### Associations of K4-A1, K6-P11 and A4-Practice scores

We found the association of A1 (I think seminars and training sessions on dairy farming are useful) was significantly different (p < 0.05) among the response of K4 (Knowledge about training and seminar) (Table 4). We found no significant difference (p >0.05) in the case of P11 (Footbath in the entrance) among K6 (Knowledge of disease spread from an outsider or neighboring farm) (Table 4). The farmers who were satisfied with their hygiene management (A4) used to have higher practice scores (PS) (Figure 2) and a significant difference (p < 0.05) in the association was found in the case of PS (Practice score) and A4 (I am satisfied about hygiene management in the farm) (Table 4).

**Table 4:**
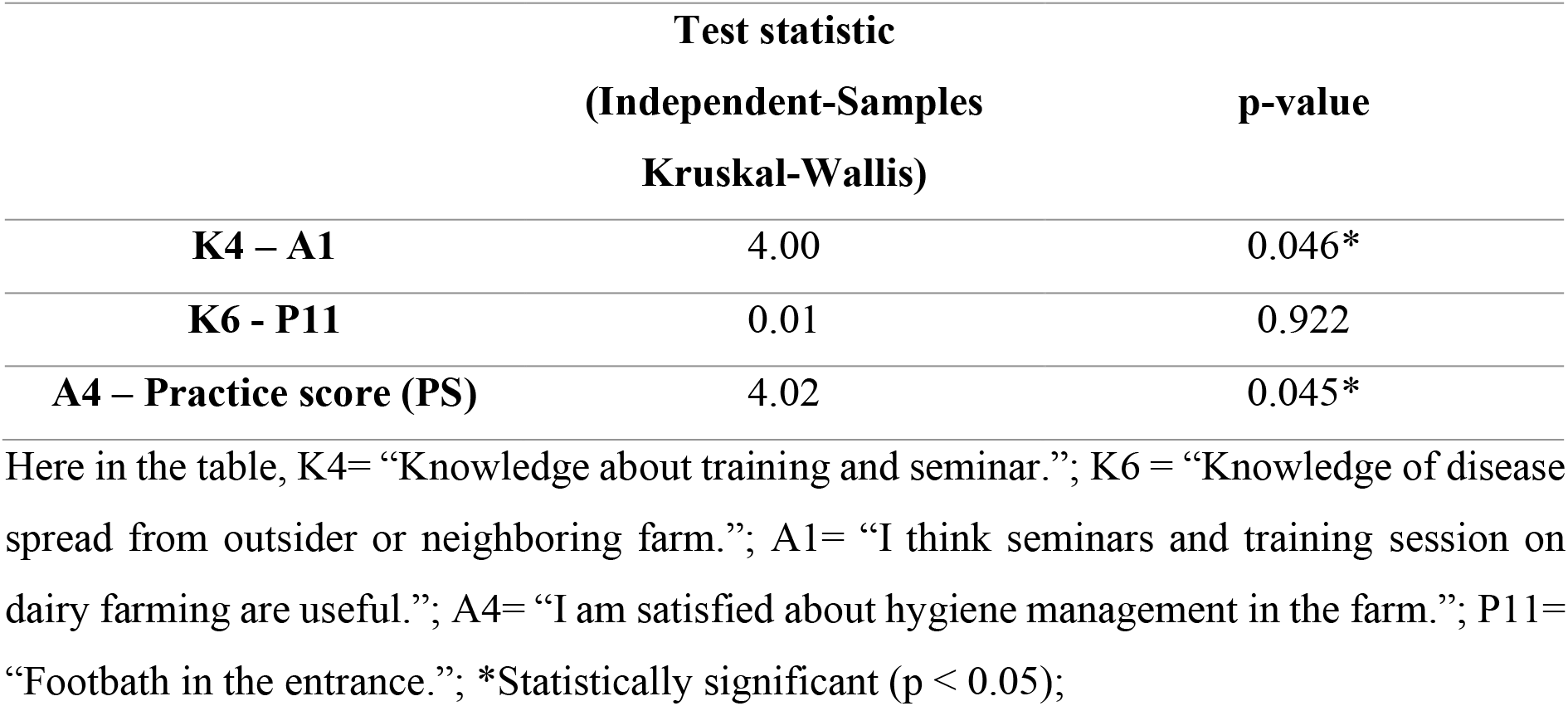
Associations between K4-A1, K6-P11 and A4-Practice score (PS).

|  | Test statistic<br>(Independent-Samples<br>Kruskal-Wallis) | p-value |
| --- | --- | --- |
| **K4 – A1** | 4.00 | 0.046* |
| **K6 - P11** | 0.01 | 0.922 |
| **A4 – Practice score (PS)** | 4.02 | 0.045* |

Here in the table, K4= “Knowledge about training and seminar.”; K6 = “Knowledge of disease spread from outsider or neighboring farm.”; A1= “I think seminars and training session on dairy farming are useful.”; A4= “I am satisfied about hygiene management in the farm.”; P11= “Footbath in the entrance.”; \*Statistically significant ( $p < 0.05$ );

**Figure 2:**
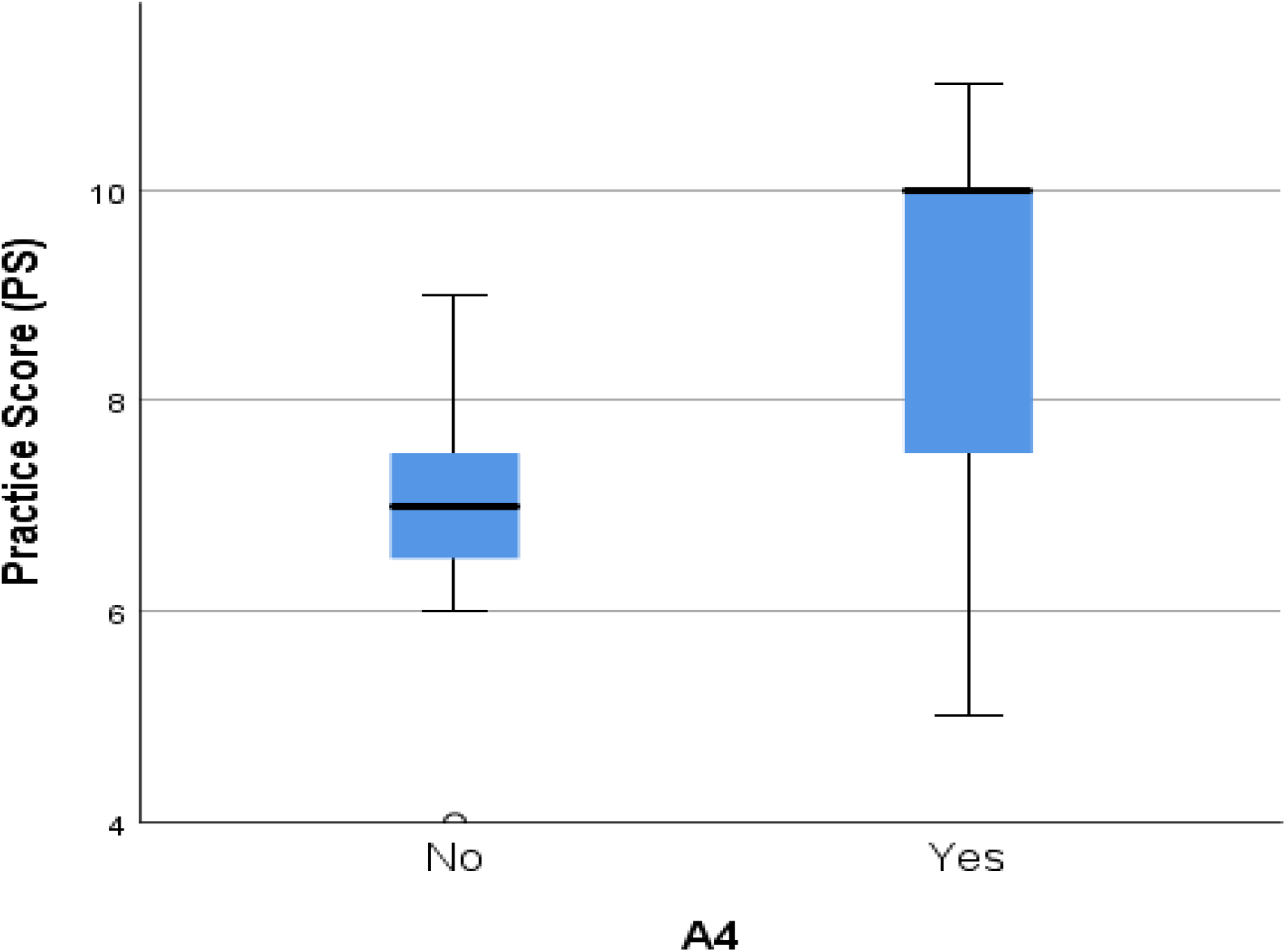
Distribution of practice scores (PS) across A4 response of the farmers (N= 15). (Here, A4= “I am satisfied about hygiene management in the farm.”)

### Correlation among knowledge, attitude and practice

Knowledge had a very strong correlation (r = 0.71) with practice and had a strong correlation (r = 0.65) with attitude (Table 5). On the other hand, attitude and practice also had a strong correlation (r = 0.64) between them (Table 5). We found significant differences (p < 0.05) in correlations between knowledge-attitude, knowledge-practice, and attitude-practice (Table 5).

**Table 5:** Correlation among farmer’s knowledge, attitude and practice regarding farm biosecurity measures.

| Variable | Spearman Correlation coefficient (r) | p-value | 95% CI lower to Upper |
| --- | --- | --- | --- |
| Knowledge – Attitude | 0.71 | 0.011* | 0.1862 to 0.8749 |
| Knowledge – Practice | 0.65 | 0.004* | 0.3001 to 0.9005 |
| Attitude – Practice | 0.64 | 0.012* | 0.1784 to 0.8729 |

\*Statistically significant ( $p < 0.05$ );

### Correlation between incidence of non-specific enteritis and biosecurity practice score

The highest non-specific enteritis incidence in 2 months was 12 experienced by the farmer and his family members, and in that particular farm, the biosecurity practice score was the lowest (4) (Table 6). The highest farm biosecurity practice score recorded was 11 and the incidence of non-specific enteritis experienced by the farmer and family members on this farm was 2 (Table 6). The lowest incidence was found 0 where the farm biosecurity score was 10 (Table 6). The correlation found between non-specific enteritis incidence and biosecurity practice score was r =-0.9232 (Figure 3). The correlation between non-specific enteritis incidence and biosecurity practice score was significant (p < 0.05) (Figure 3).

**Table 6:**
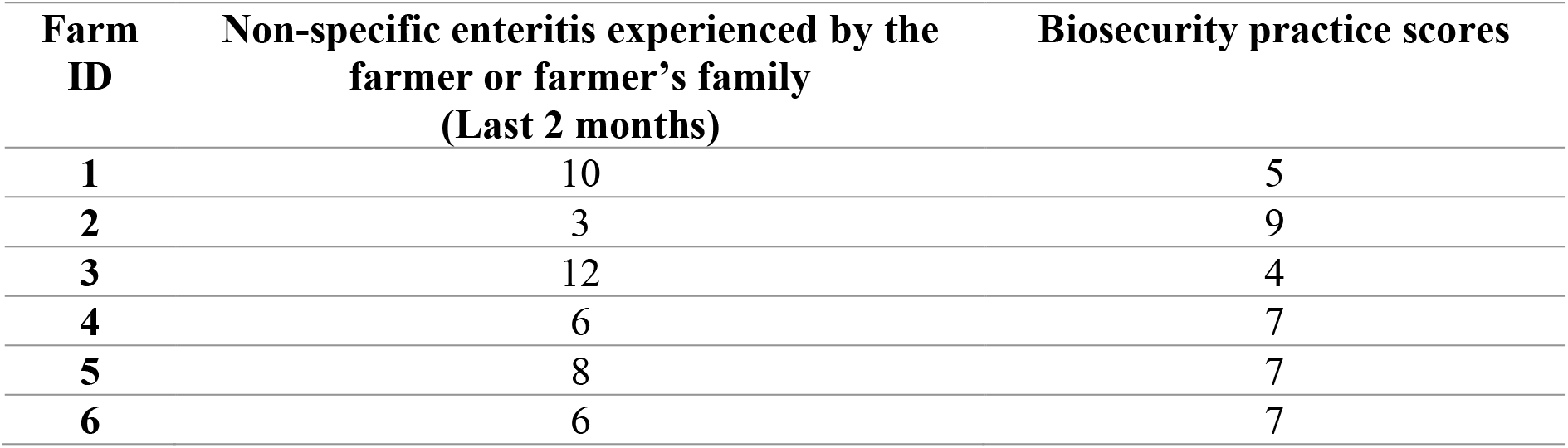
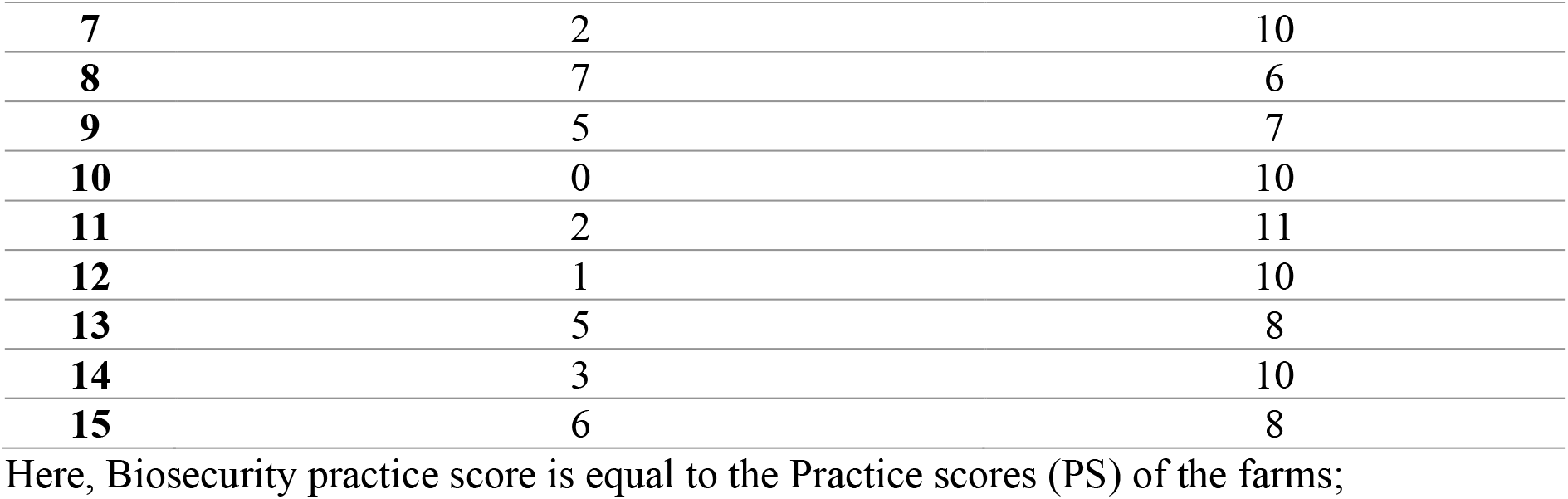
Biosecurity practice scores and the number of non-specific enteritis experienced by the farmers (N=15) or family members.

| Farm ID | Non-specific enteritis experienced by the farmer or farmer's family (Last 2 months) | Biosecurity practice scores |
| --- | --- | --- |
| 1 | 10 | 5 |
| 2 | 3 | 9 |
| 3 | 12 | 4 |
| 4 | 6 | 7 |
| 5 | 8 | 7 |
| 6 | 6 | 7 |
| 7 | 2 | 10 |
| 8 | 7 | 6 |
| 9 | 5 | 7 |
| 10 | 0 | 10 |
| 11 | 2 | 11 |
| 12 | 1 | 10 |
| 13 | 5 | 8 |
| 14 | 3 | 10 |
| 15 | 6 | 8 |

Here, Biosecurity practice score is equal to the Practice scores (PS) of the farms;

**Figure 3:**
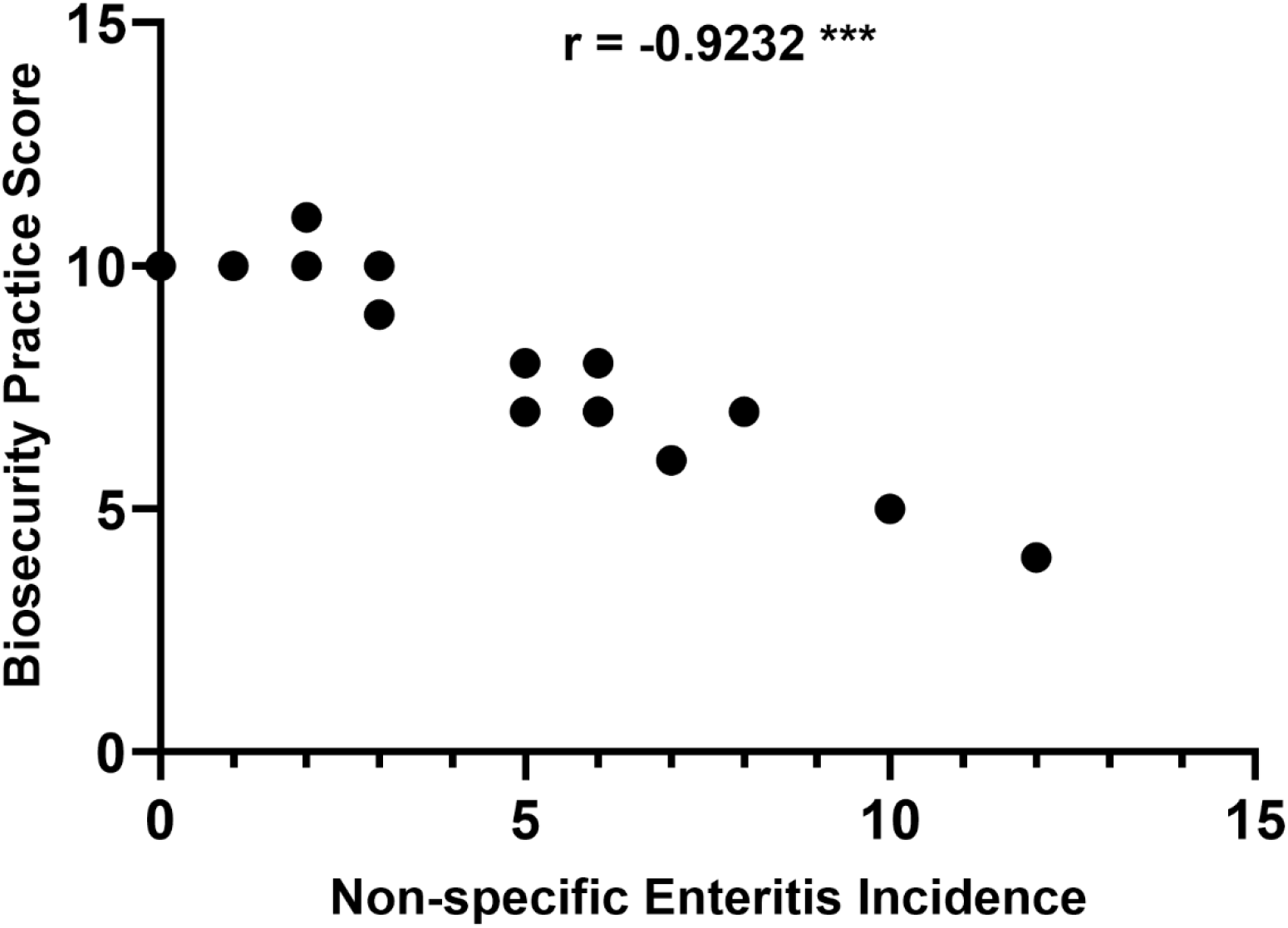
Distribution and correlation of non-specific enteritis incidence and biosecurity practice scores of the farmers. (Here, ‘r’ = Spearman’s correlation coefficient; Significance level was considered as p-value <0.05; ‘***’= p-value <0.001;)

## Discussion

In this current study, a relatively small sample size was used, that is because the study area does not accommodate many commercial dairy farms and also all farms cannot be visited due to lack of time and unwillingness of the owner to participate in the survey. However, the present study contains enough information to understand the knowledge, attitude, and practice of biosecurity among the farmers of the study area.

Previous studies show that knowledge, attitude, and practice are associated with one another, and demographic characteristics can also have an impact on knowledge, attitude, and practice (Can & Altuğ, 2014; Jafari-Gh et al., 2020; Makita et al., 2020; Mateo et al., 2021). So, we hypothesized – (1) Demographic characteristics have influences on the knowledge, attitude, and practice of biosecurity measures; (2) Knowledge, attitude, and practice of biosecurity measures are strongly correlated. Furthermore, animal and human health are closely related because of the possibility of direct or indirect pathogen transmission between them, and dairy farms and dairy products are considered to be possible sources of pathogens that could cause human health problems including gastroenteritis (An et al., 2018; Pell, 1997; Stein & Katz, 2017; Youssef et al., 2021). So, without proper biosecurity measurements, there is a possibility of transmission of pathogens from the dairy farm environment or dairy products to the farmers or dairy product consumers. Hence, we developed the hypothesis-there is a strong correlation between the practice of biosecurity measures and the incidence of non-specific enteritis in farmers and their family members who are directly or indirectly related to the farms or consume milk from that farm.

### Frequency percentages and mean scores of knowledge, attitude and practice

Seminars and training sessions about biosecurity can increase awareness and 80% of the farmers had knowledge about seminars and training sessions (K4) (Table 1). But only 53.3% agreed that seminars and training sessions are useful (A1) (Table 1). Another impactful measure is to keep disease records on farms but only 60% of the farmers knew about record-keeping (K5) (Table 1) contrast to the European survey which reported that about 73% to 91% of dairy farmers used to keep a record (Denis-Robichaud et al., 2019). The disease can be spread through indirect or direct contact in various ways like different farm visiting personnel and different equipment from other farms (Brennan & Christley, 2012). So, it is necessary to minimize the risk by adopting biosecurity measures but less than half of the farmers (46.7%) knew about the possible spreading of diseases from outsider’s entrance or neighboring farms (K6) (Table 1) and only 13.3% used to have footbath on the entrance of the farm (P11) (Table 2). However, 73.3% believed that only necessary visits should be allowed in the farm (A5) (Table 1). Regular cleaning and disinfection reduce biosecurity risks of a farm. In present study, 93.3% of the farmers did regularly cleaning and disinfection of their farms (P10) (Table 2). Another aspect of reducing disease risk is to test diseases before buying any new animals and vaccination against diseases (Denis-Robichaud et al., 2019). According to a previous study, about 86% of Irish dairy farmers and 70% of Canadian dairy farmers used to vaccinate their animals with at least a single dose (Denis-Robichaud et al., 2019). In the present study, 86.7% practiced vaccination of animals against common contagious diseases (P7) (Table 2). A previous study also reported that around 46% of Canadian farmers used to test for disease before introducing new animals to the farm (Denis-Robichaud et al., 2019). However, in the current study most farmers (86.7%) used to test diseases before buying new animals (P1) (Table 2).

### Comparison of knowledge, attitude and practice scores of demographic characteristics

From (Table 3), D1 (Farmer’s age) shows that elderly farmers (>40 years) tend to have more knowledge (4.0 KS) and have better practice (9.3 PS). Maybe the elderly farmers (>40 years) are more likely to gather information and implement biosecurity practices by replicating practices from other farms but less likely to believe that these practices are actually necessary. However, we found no significant differences (p > 0.05) in KS, AS, and PS among the age groups which is supported by the previous findings (Can & Altuğ, 2014). D2 (Farmer’s educational qualification) demonstrates that farmers with a graduation level of educational background tend to adopt better biosecurity practices (8.6 PS) but their attitude and knowledge regarding biosecurity may lack (Table 3). A previous study found that highly educated farmers tend to have better biosecurity scores (Can & Altuğ, 2014). It was also found that educational level had a significant impact on farmers’ knowledge, attitude, and practice (Jafari-Gh et al., 2020). However, we did not find any significant differences (p >0.05) in KS, AS, and PS among educational level (D2) which contradicts the findings of (Can & Altuğ, 2014) (Table 3).

D3 (Farmer’s farming experience) shows that farmers with less experience (< 10 years) had better knowledge (3.7 KS) but those who had experienced over 20 years had better attitude (4.5 AS) and practice (8.5) (Table 3). It is possibly because; the less experienced farmers try to thrive knowledge for the betterment of the farm but cannot implement the knowledge. No significant differences were found (p > 0.05) among farming experience (D3) which is supported by the findings of (Can & Altuğ, 2014) (p > 0.05). In the case of income class (D4), results depict that, farmers with higher income (> $500 / month) had lesser knowledge about biosecurity but a better attitude (3.6 AS) and practice (8.1 PS) than the farmers of middle-income ($250-500/ month) group (Table 3). However, the differences of KS, AS, and PS among income class were not significant (p > 0.05) (Table 3), but this result indicates that having knowledge about biosecurity does not always results in practices of biosecurity measures. As previously noted by veterinarians, a lack of knowledge of biosecurity is not only the reason for implementing biosecurity in farms but also farmers’ attitudes and will also play important roles (Pritchard, Wapenaar, & Brennan, 2015). A previous study also found that higher income resulted in higher biosecurity scores (Can & Altuğ, 2014). It was also reported that higher income has a high impact on the knowledge, attitude, and practice of a farmer (Jafari-Gh et al., 2020).

From current findings, D5 (Age of the farm) shows that farmers from the farms which existed for 3 to 5 years had better knowledge (4.5 KS), attitude (4.0 AS), and practice (8.6 PS) (Table 3). This depicts that certain periods after the starting of farms perform better in biosecurity measures but in the state introductory period the farmers may lack resources to access information about biosecurity measures. On the other hand, farmers from farms with ages more than 5 years may be reluctant to consider biosecurity measures as necessary because the farm has already survived a long time. D6 (Number of animals in farm) shows that farms with less than 15 animals have better knowledge (4.3 KS) and attitude (4 AS) but farms with more than 25 animals have better practice (8.6 PS) (Table 3). Larger herd size results in higher biosecurity scores were also found in a previous study (Can & Altuğ, 2014). Also, farms with large herd sizes may have better biosecurity because these farms have a higher risk of losses due to diseases (Jafari-Gh et al., 2020). However, we didn’t find any significant differences (p >0.05) of KS, AS, and PS in the case of D5 and D6 (Table 3). Finally based on the current study findings, we rejected our alternative hypothesis that demographic characteristics have influences on the knowledge, attitude, and practice of biosecurity measures.

### Associations of K4-A1, K6-P11 and A4-Practice scores

Believing seminars and training sessions could be useful (A1) was significantly different (p < 0.05) between the farmers who had knowledge of training and seminars (K4) and who didn’t (Table 4). Though training sessions and seminars are important for improving biosecurity and policy making of farms, negative attitudes and fatigue still exist among farmers. To improve the situation, the responsible factors should be identified and alternative approaches need to be formulated to motivate and engage the farmers in seminars and training (Hamilton, Evans, & Allcock, 2019). In this present study, whether farmers knew about the risk of disease spreading through neighboring farms or outsiders (K6), didn’t significantly affect the practice of using footbath (P11) (p > 0.05) (Table 4). But using footbaths on the farm can improve the bovine feet health and reduce biosecurity risk in farms (Fjeldaas et al., 2014). Additionally, using footbath in farms should be an essential practice in the current study area as it is considered to be a hot spot for contagious diseases like FMD (Rahman et al., 2020). The current study results also revealed that practicing better biosecurity measures (higher practice score) was closely related to having satisfaction with the hygiene management of the farm (A4) (Table 4; Figure 2). But that doesn’t exclude the chances that farmers will not be satisfied with less biosecurity practices. Hence, if it could be possible to broaden the satisfaction margin of the farmers then they would be automatically encouraged to adopt better biosecurity measures.

### Correlation among knowledge, attitude and practice

Knowledge, attitude, and practice had a strong positive correlation with one another (Table 5). That means a change in one of these variables will affect another factor in a positive direction. If the farmer had better knowledge of biosecurity, it would result in a positive attitude toward biosecurity measures and better practices. However, farmers’ perceptions of biosecurity may evolve and change and may not be consistent over time (Brennan & Christley, 2013). So, knowledge of biosecurity should be disseminated with a standard guideline, and regular training should be provided to keep the farmers updated with new information. Previous research has also found that improved knowledge leads to more positive attitudes, and positive attitudes lead to more biosecurity practices (Makita et al., 2020). So, three of these factors coexist together for the improvement of biosecurity measures in farms. Hence, based on current study findings (Table 5), we accepted our alternative hypothesis that there are associations among the knowledge, attitude, and practice of biosecurity measures.

### Correlation between incidence of non-specific enteritis and biosecurity practice score

The notion of biosecurity has gained importance over the years due to the numerous hazards and heightened animal-associated risks brought on by demographic and environmental changes, along with globalization and international exchange (Lytras, Xia, Hughes, Jiang, & Robertson, 2021). Dairy cattle had been identified as potential reservoir pathogens such as *Campylobacter* which causes human gastroenteritis (An et al., 2018). Gastroenteritis-causing pathogens like *Escherichia coli* had also been identified in dairy milk and farms as well (Stein & Katz, 2017). These pathogens can easily transmit to humans via milk or direct contact with farm utensils due to a lack of biosecurity measures. The strong correlation (r = -0.9232) between the incidence of non-specific enteritis and farm biosecurity practice score found in the current study (Figure 3) demonstrates that adaption of more biosecurity measures reduces the incidences of non-specific enteritis among farmers and their family members who are directly or indirectly related to the farms or consume milk from that farm. Previous study shows that the implementation of good biosecurity measures reduces the transmission of pathogens from livestock to human (Youssef et al., 2021). Moreover, limiting dairy farms as the only reason for non-specific enteritis would not be a wise discussion. Because enteritis could also develop from other food sources such as broiler meat (la Mora et al., 2020). But the significant (p < 0.05) correlation between enteritis and farm biosecurity practice score found in the current study cannot be ignored as well (Figure 3). However, the finding in our current study about the correlation of enteritis and biosecurity measures do not claim that the incidences of non-specific enteritis only depend on the biosecurity measures of the firm, rather from our findings, the assumption may be made that biosecurity practice does influence the health of the individuals who are directly or indirectly connected to the products or the environment of the farms. For a stronger claim on the biosecurity practice-enteritis relationship, a thorough study would be needed for identifying enteritis-causing organisms in the farm environment or farm products, and the causal agent of enteritis in the individuals who are in contact with the firm, and analysis of genetic homology of those microorganisms.

The current study revealed that demographic characteristics do not influence knowledge, attitude, and practice of biosecurity measures. Knowledge, attitude, and practice are highly and positively correlated with one another. With better knowledge, the farmers’ attitude and practice of biosecurity measures improve. Biosecurity score is also correlated with non-specific enteritis incidence. Having higher farm biosecurity practice measures reduces the incidence of non-specific enteritis in individuals who are directly in contact with the farm or consume milk from that farm. However, awareness is needed to be increased for a better understanding and implementation of biosecurity measures. Finally, further studies are needed to establish a strong claim.

## Conclusion

Our study reveals most of the small-scale dairy farmers of Sylhet District, Bangladesh, are experiencing non-specific enteritis. And the knowledge, attitudes, and current biosecurity practices are yet to gain a satisfactory level to prevent zoonosis such as non-specific enteritis. So, the farmers need more awareness and relevant training to enhance their biosecurity practices regarding public health importance.

## Acknowledgements

We like to disclose our gratitude to the Chairman of Dairy Science department to assist us with this study.

